# Functional dissociation of the language network and other cognition in early childhood

**DOI:** 10.1101/2022.08.11.503597

**Authors:** K.J. Hiersche, E. Schettini, J. Li, Z.M. Saygin

## Abstract

Is language distinct from other cognition during development? Does neural machinery for language emerge from general-purpose neural mechanisms, becoming tuned for language after years of experience and maturation? Answering these questions will shed light on the origins of domain-specificity in the brain. We address these questions using precision fMRI and found young children (35 months-9 years) show domain-specific, left-lateralized language activation, and the language network is not responsive to domain-general cognitive load. Additionally, the cortically adjacent multiple demand network is selective to cognitive load, but not to language. These networks show higher within vs. between-network functional connectivity. This connectivity is stable across ages (examined cross-sectionally and longitudinally), whereas language responses increase with age and across-time within subject, reflecting a domain-specific developmental change. Overall, these findings suggest that domain-specificity, even for uniquely human cognition like language, develops early and distinctly from mechanisms that presumably support other human cognition.

**Significance Statement:** This study provides evidence of a double dissociation between neural processors for language and domain-general cognition (specifically, cognitive load) in young children. These results refute previous claims that linguistic function emerges from domain-general mechanisms supporting non-linguistic, higher-order cognition, and with both cross-sectional and longitudinal analyses, show that the continued development of linguistic specialization is also unrelated to domain-general mechanisms that support cognitive flexibility and executive function. This work emphasizes the importance of single-subject fMRI analyses with multiple tasks to explicitly dissociate cognitive constructs, and the importance of longitudinal research for scientific rigor, even in toddlers and preschoolers who are difficult to image with fMRI.

## Introduction

A domain-specific network of brain regions support language, a uniquely human cognitive skill; remarkably, language develops naturally through exposure, without requiring explicit training or formal education in the first few years of life (Lenneberg, 1967; Sharp & Hillenbrand, 2008). How does a child’s brain support this early language acquisition? Perhaps language skills develop via the same domain-general cognitive processes that support other cognitive skills, and therefore, the language network may initially show sensitivity to other, more general cognitive skills. Alternatively, young children’s language networks may show early specialization for language, suggesting domain-specificity even for uniquely human cognition like language.

Prior work in adults demonstrates that high-level language is dissociable from other cognitive processes. Language function is supported by a domain-specific, left-lateralized, frontotemporal language network (Fedorenko et al., 2010, 2011; Ojemann, 1991; Vigneau et al., 2006), that is *selective* for the semantic and syntactic properties of language (Fedorenko et al., 2010). The language network is domain *specific*; it does not respond to other higher-order cognitive skills. These skills, like performing arithmetic, cognitive control, or working memory, are recruited by the cortically adjacent, domain-general fronto-parietal multiple demand (MD) network (Duncan, 2010; Fedorenko et al., 2013). The MD network responds to a variety of tasks when contrasting a more cognitively demanding condition with a less demanding condition (Hard vs. Easy task effect; cognitive load). In adults, these two networks are distinct in both their function and connections (Blank et al., 2014; Fedorenko et al., 2011, 2012). Similar patterns of activation and connectivity profiles during development would suggest that language is distinct from other cognition, or at least recruits different neural mechanisms. Is the language network already dissociated from domain-general cortex in young children who are still developing both domain-specific language and domain-general cognitive abilities?

If these two networks are not dissociable, but instead entangled early in development, it may suggest that the origins of domain-specificity in the brain are a result of early scaffolding by generalized mechanisms (Karmiloff-Smith, 2015). Perhaps the mechanisms necessary for the brain to gain increasing specialization for processing language are not so different from other types of complex mental operations (e.g., holding items in memory (Hamrick et al., 2018; Ullman, 2004) or performing arithmetic) which recruit domain-general cognitive processes with increasing task demand, or perhaps executive control may help bootstrap language learning by supporting alternative processing mechanisms or management strategies (Edgin et al., 2015; Saffran & Thiessen, 2007). Indeed, some behavioral work suggests that domain-general skills, such as attention, play a role in typical language development (Miller, 2006; West et al., 2021), but this has not been fully explored in the developing brain. No prior work has yet explored this question of double dissociation of language and domain-general cortex in young children and their continued development.

Previous studies have investigated the development language network on its own in children using task-based fMRI; however, results are discordant. Language network development in school-age and older children show activation in traditional perisylvian language regions (Enge et al., 2020; Weiss-Croft & Baldeweg, 2015) with increasing activation in the left hemisphere (LH) with age (Weiss-Croft & Baldeweg, 2015), but it is unclear whether temporal language regions emerge first (during early childhood at ages 5-7) (Berl et al., 2014; Weiss et al., 2018), and frontal regions develop later (after 7-10 years of age, Berl et al., 2014; Wang et al., 2021); or if both frontal and temporal language network activation emerges in early childhood, with a systematic decline in right hemisphere (RH) language selectivity with age (Olulade et al., 2020).

These discordant results may be due to confounds of task diffulty and/or lack of spatial precision when defining the language network. Task performance, particularly in expressive language studies, may confound results due to age-related general effects on performance. Additionally, given that the precise location of these language regions can vary across subjects (Amunts et al., 1999; Juch et al., 2005; Tomaiuolo et al., 1999), the boundaries between language and cortically adjacent domain-general cortex may have been blurred in previous work using group-level regional masks, leading to uncertainty about whether language cortex is continuing to develop. Therefore, it is important to either account for task performance during expressive tasks or use a passive language task for children, with stimuli that are semantically meaningful and have rich syntactic structure. Additionally, it is important to use subject-specific functional regions of interest (as demonstrated in e,g., Fedorenko et al., 2012) that account for between subject variability in functional organization (Gratton et al., 2018) when examining the dissociation between language and spatially-adjacent domain-general cortex.

Identifying the developmental dissociation of language cortex from domain-general regions will help us better understand the origins of the neural mechanisms that underlie unique human knowledge. If language cortex initially shows little selectivity to linguistic content but engages in other cognition such as working memory and continues to refine over time (e.g., becoming less engaged in working memory while gaining linguistic selectivity), then domain-general processes may bootstrap the development of uniquely human cognitive skills like language (i.e., the language system may emerge through the same domain-general skills applied in other contexts; McMullen & Saffran, 2004; Rakison & Yermolayeva, 2011; Saffran & Thiessen, 2007). Alternatively, domain-general cortex may represent multiple domains (i.e., language, working memory, cognitive demand, etc.) early in development, so that it can serve a general purpose during adulthood (i.e., MD cortex in young children would support both working memory and linguistic content). In either case, it would be of interest to know whether the mechanisms that support these two seemingly discrete processes overlap during childhood. If we observe that the language network is domain-specific (selective only for linguistic processing but not cognitive load), the MD network is domain-general (selective to cognitive load but not responsive to linguistic properties), that these networks are also differentiated in their connectivity, and that they develop independently of one another, this would support a third hypothesis: there is a (possibly) innate proto-organization of domain-specificity in the brain.

Here we investigate the development of the canonical high-level language network and its differentiation from domain-general cortex in a group of young children ages 35 months to 9 years of age. We leverage individual subject data on two separate fMRI tasks, use single-subject fROIs defined in native anatomy, to account for variability in the precise location of these regions (Fedorenko et al., 2010, 2013), and directly compare linguistic and nonlinguistic activation in the same subjects. We also examine the within- and between-network resting-state connectivity of the language vs. MD fROIs. We assess any age-related changes in linguistic selectivity and connectivity both cross-sectionally and longitudinally in a subset of participants.**Results**

### Is the high-level language network already selective for linguistic content in young children?

To visualize overlap of language activation in our sample (see three example subjects in **Figure 1A)**, we first created probabilistic atlases (**Figure 1B**). We defined language activation using the contrast of meaningful Sentences > Nonsense sentences (Sn>Ns) which defines high-level language cortex by controlling for the rhythm and prosody of human speech. This probabilistic atlas revealed language activation across specific regions within left-hemispheric temporal and frontal cortices. While there was large variability across individuals (highlighting the need for individual-subject definitions of fROIs in this literature), there was still substantial overlap of language activation within the left temporal regions, and moderate overlap in left frontal regions, across subjects. Right-hemispheric activation was less prominent, with modest overlapping activation in the right temporal regions and no right frontal regions survived thresholding. Language activation fell within the expected frontotemporal regions suggesting that the child language network falls largely within search spaces (black outlines, **Figure 1A**) defined and used in adult studies of the language network.

**Figure 1.**
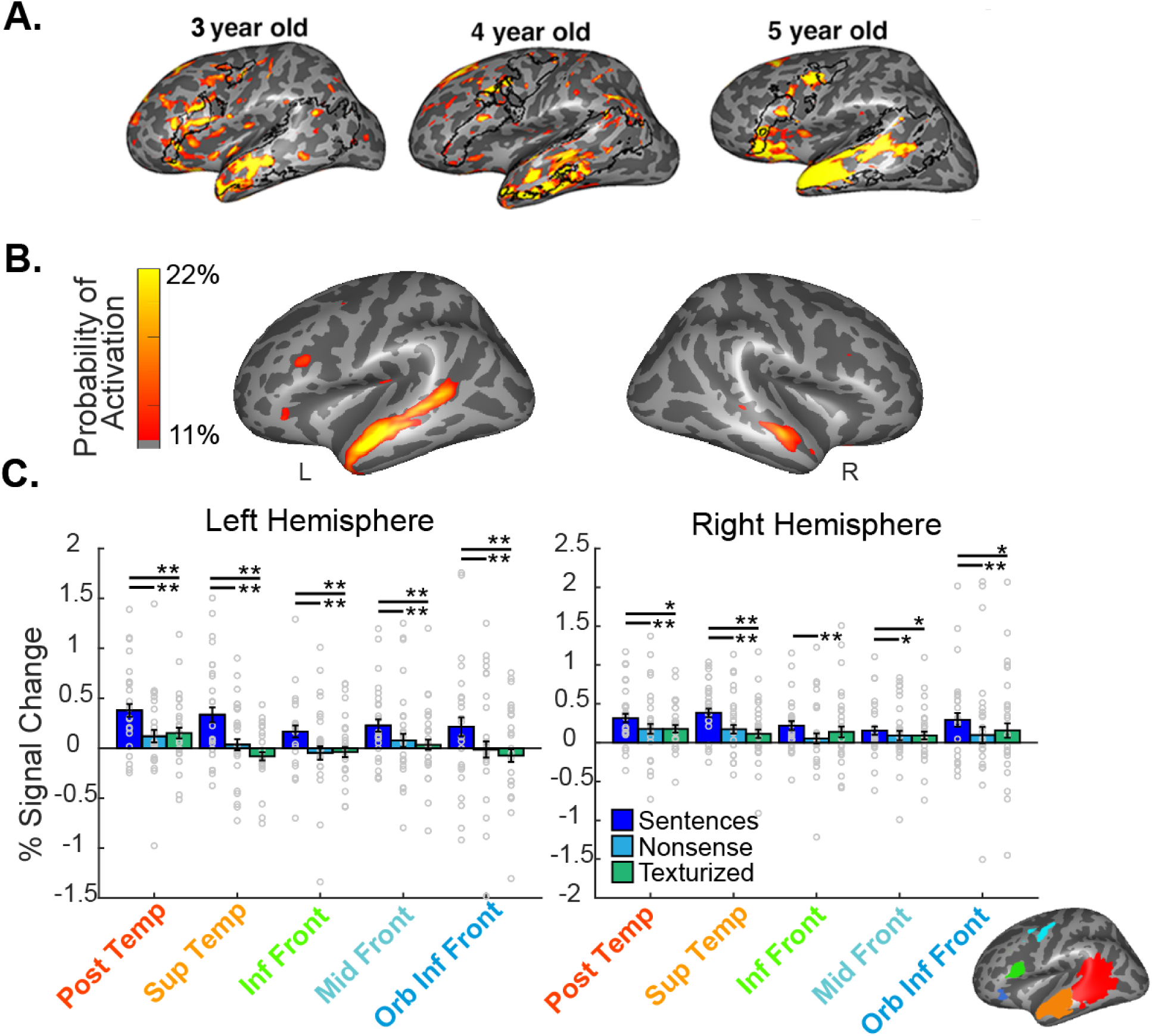
Selectivity of the high-level language network. A. Activation to Sn>Ns in the left hemisphere for example subjects (thresholded at P_uncorrected_ < 0.005). Black outlines show search spaces for the five language regions registered to the example participant brain and used for creating fROIs. B. Probabilistic atlas showing common activation (Sentences>Nonsense sentences) during both runs of the language task (z > 2.58) in at least 5 (11%) participants and a maximum of 10 (22%) participants. C. Mean percent signal change to all language task conditions in whole sample (N=45). Inset shows parcel search spaces used for created fROIs. Asterisks indicate level of significance (two-tailed, paired t-test, *p<0.05, **Bonferroni-Holm p < 0.05, corrected for 10 comparisons within hemisphere). Error bars denote standard error. Individual subject data points shown with hollow grey circles.

Next, we quantitatively examined the selectivity of the language network. We compared the responses of subject-specific language fROIs (5 bilateral regions) to each condition of the language task (meaningful Sentences, meaningless Nonsense sentences, and Texturized sound controlling for low-level auditory features) using a repeated-measures ANOVA (rmANOVA) of condition by fROI. For all rmANOVAs, Greenhouse-Geisser corrected p-values are listed when appropriate; uncorrected p-values are listed for post-hoc comparisons and survive Bonferroni-Holm multiple-comparison correction unless otherwise noted. In the LH, we observed a significant main effect of fROI (F(4,164)=7.11, p=2.66×10^-5^, η^2^=0.028), condition (F(2,82)=12.49, p=1.84×10^-5^, η^2^=0.064) and a significant fROI by condition interaction (F(5.57,228.57)=2.86, p=0.012, η^2^=0.007), such that all LH fROIs were significantly more responsive to Sentences than both control conditions (vs Ns all p<0.05, and vs. Textured sound or Tx: all p<0.05, as revealed by follow-up paired t-tests, see **Figure 1C**, **Table 1**). In the right hemisphere (RH), we again see a significant main effect of condition (F(2,76)=8.90, p=3.39×10^-4^, η^2^=0.062; no significant fROI effect: F(3.17,120.27)=2.56, p=0.055, η^2^=0.018) and an fROI by condition interaction (F(5.17,196.55)=3.16, p=0.0080, η^2^=0.009), such that all RH fROIs were significantly more responsive to Sentences than Nonsense and Textured Sound (except the right inferior frontal, most comparisons surviving correction, see **Table 1**). While we observe selectivity to linguistic content in fROIs of both hemispheres, given the language network is characteristically left-lateralized, next we formally compared language selectivity across hemispheres. We found a main effect of both fROI (F(2.82,106.98)=4.12, p=0.010, η^2^=0.023) and hemisphere (F(1,38)=5.77, p=0.021, η^2^=0.014; no significant interaction: F(3.14,119.31)=0.54, p=0.66, η^2^=0.002), such that selectivity of the LH fROIs were significantly higher than the RH (averaged selectivity across fROIs: t(38)=2.40, p=0.021, *d*=0.38). Overall, we find evidence of a left-lateralized language network that is selective to linguistic content in a young sample of children.

**Table 1:** Selectivity of language network fROIs. Two-tailed, paired t-tests comparing the Sentences condition to control conditions (Nonsense sentences and Texturized sound), following significant condition by fROI interaction in rmANOVA. T value, significance, and Cohen’s D effect sizes listed. *p < 0.05, **Bonferroni-Holm p < 0.05, corrected for 10 comparisons within hemisphere).

|  | *Sentences vs. Nonsense* |  |  | *Sentences vs. Texturized* |  |  |
| --- | --- | --- | --- | --- | --- | --- |
| *LH* | *t(df)* | *p* | *d* | *t(df)* | *p* | *d* |
| Posterior Temporal | $t(41)=4.98$ | $1.12 \times 10^{-5**}$ | 0.77 | $t(41)=3.31$ | $0.003**$ | 0.48 |
| Superior Temporal | $t(41)=4.74$ | $2.54 \times 10^{-5**}$ | 0.73 | $t(41)=5.17$ | $1.80 \times 10^{-6**}$ | 0.86 |
| Inferior Frontal | $t(41)=2.85$ | $0.007**$ | 0.44 | $t(41)=3.53$ | $0.001**$ | 0.54 |
| Middle Frontal | $t(41)=2.66$ | $0.011**$ | 0.41 | $t(41)=3.88$ | $3.72 \times 10^{-4**}$ | 0.60 |
| Orbital Inferior Frontal | $t(41)=2.40$ | $0.021**$ | 0.37 | $t(41)=2.72$ | $0.010**$ | 0.42 |
| *RH* | *t(df)* | *p* | *d* | *t(df)* | *p* | *d* |
| Posterior Temporal | $t(38)=3.45$ | $0.0010**$ | 0.55 | $t(38)=2.29$ | $0.028*$ | 0.37 |
| Superior Temporal | $t(38)=5.15$ | $8.26 \times 10^{-6**}$ | 0.82 | $t(38)=5.97$ | $6.16 \times 10^{-7**}$ | 0.96 |
| Inferior Frontal | $t(38)=3.28$ | $0.0020**$ | 0.52 | $t(38)=1.93$ | $0.061$ | 0.31 |
| Middle Frontal | $t(38)=2.09$ | $0.043*$ | 0.34 | $t(38)=2.11$ | $0.041*$ | 0.34 |
| Orbital Inferior Frontal | $t(38)=3.35$ | $0.0020**$ | 0.54 | $t(38)=2.34$ | $0.023*$ | 0.38 |

### Does the selectivity of the language network change with age?

To examine age-related changes in language selectivity, we repeated the rmANOVA of experimental conditions by fROI, adding age as a covariate (rmANCOVA). Replicating the results from above, we found a main effect of condition (LH: F(3.35, 133.97)=7.21, p=8.52×10^-5^, η^2^=0.030), fROI (LH: F(2,80)=13.85, p=6.86×10^-6^, η^2^=0.068), and condition by fROI interaction (LH: F(5.52,220.64)=2.84, p=0.013, η^2^=0.007), but importantly we also observed an age by condition interaction (LH: F(2,80)=5.45, p=0.006, η^2^=0.028). Post-hoc analyses showed that the LH fROIs responses increase with age specifically for the Sentences condition (averaged PSC across fROIs; r(40)=0.40, p=0.0090; partial correlation controlling for motion: r(40)=0.40, p=0.010), but not in responses to either control condition (Ns: r(40)= 0.081, p=0.61, partial: r(40)=0.038, p=0.811; Tx: r(40)=-0.0086, p=0.96, partial: r(40)=0.0018, p=0.99). We did not see a significant age by condition effect in the RH (F(2,74)=0.67, p=0.52, η^2^=0.005). Therefore, while a left-lateralized language cortex is already present in this young sample, the left language regions continued to gain language selectivity with age.

A subset of participants was also scanned longitudinally and therefore completed the language task at multiple timepoints (N=23) approximately one and a half years apart. We examined changes in selectivity of the language network across time within each child (reliable portion of language fROIs at timepoint two were registered back to subject space at timepoint one, see **Methods**). In a rmANOVA of condition by timepoint by fROI, we found significant main effects of condition (LH: F(2,32)=4.33, p=0.022, η^2^=0.050) and fROI (LH: F(4,64)=4.61, p=0.0020, η^2^=0.047), as well as a condition by fROI interaction (LH: F(8,128)=3.77, p=5.40×10^-4^, η^2^=0.010), in the LH. Most importantly, we found a significant condition by timepoint interaction (LH: F(2,32)=4.30, p=0.022, η^2^=0.02), such that only the Sn condition significantly increased across timepoints within subjects (post-hoc two-tailed, paired-samples t-tests: Sn: t(16)=3.07, p=0.007), with no significant change in either control condition (Ns: r(16)=-0.26, p=0.80, Tx: r(16)=0.65, p=0.53), aligning with our cross-sectional results. On the RH, however, while we found a significant main effect of fROI (F(4,60)=5.86, p=4.78×10^-4^, η^2^=0.081) and condition by fROI interaction (F(4.28,64.20)=3.44, p=0.011 η^2^=0.012), there was no significant timepoint by condition interaction (F(2,74)=0.67, p=0.52, η^2^=0.005) or main effect of condition (F(1.45,21.77)=2.33, p=0.13, η2= 0.019). Taken together, both cross-sectional and longitudinal results suggest that the age-dependent increases in LH language selectivity were likely due to an increased response to linguistic content rather than a decreased response to control conditions.

To ensure differences in language selectivity across time were not due to motion, we compared the mean framewise displacement within subject, across time. Participant motion was not significantly different from timepoint 1 to timepoint 2, (two-tailed, paired samples t-test, t(22)=1.81, p=0.085, d=0.38); therefore, it is unlikely that motion explains the increase to linguistic content over time.

### Is the high-level language network specific for linguistic content and differentiated from domain-general cortex?

After demonstrating selectivity of the language network of young children, we examined the *functional specificity* of these early-developed language regions by comparing the same children’s activation to a spatial working memory task’s Hard and Easy conditions (cognitive load, i.e., the Hard condition elicits higher engagement of the MD network, tapping the domain-general processes (Fedorenko et al., 2013; Schettini et al., 2023). Activation maps for an example participant illustrate the distinct responses for these two tasks, in which language activation (Sentences > Nonsense) falls largely within the language parcel search spaces and the working memory task (Hard > Easy) activates regions outside of this network, particularly in the frontal and parietal lobes (**Figure 2A**).

**Figure 2:**
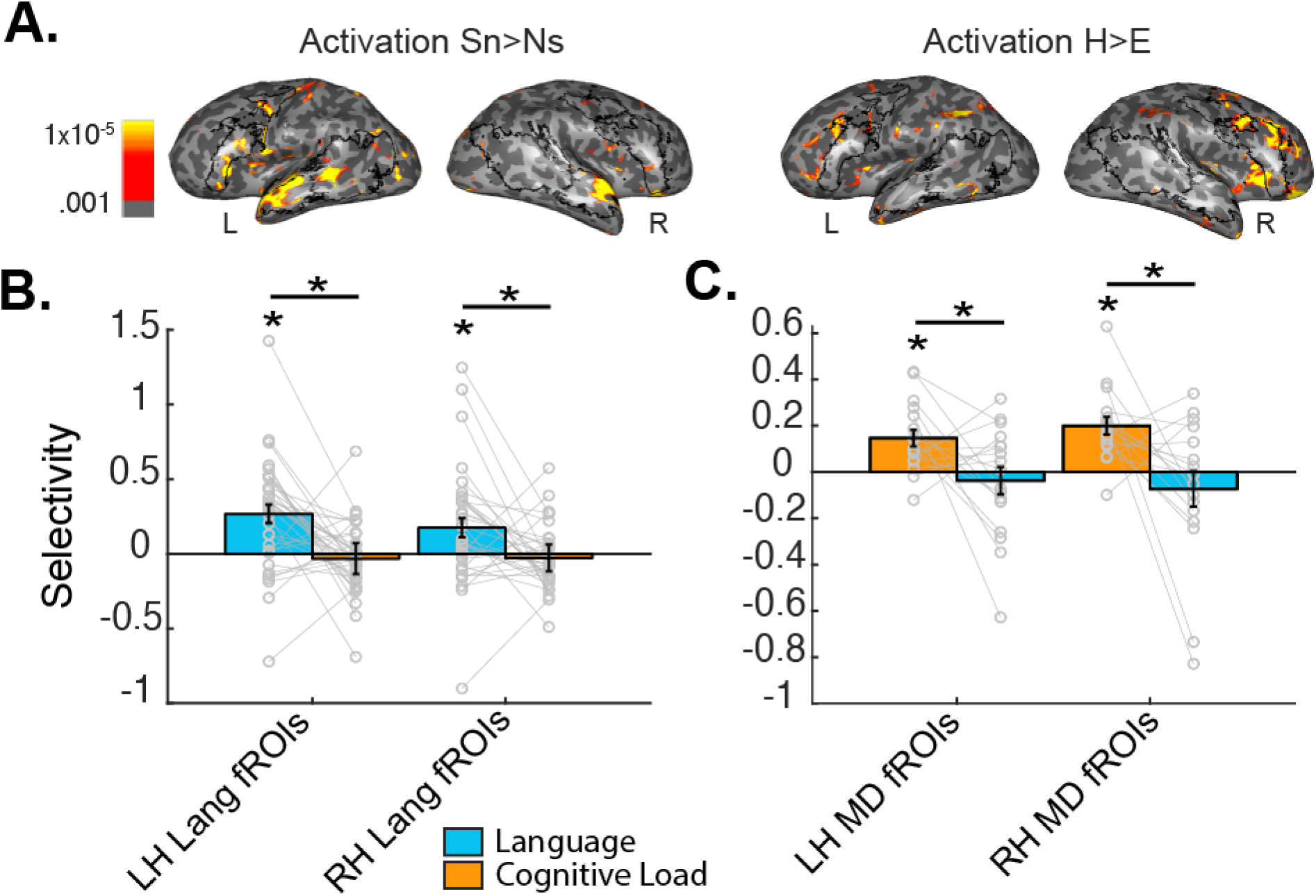
Language and cognitive load activation in language and MD fROIs. A. Language (left) and cognitive load (H>E, working memory task) (right) activation for an example subject (age 7.8 years, threshold at P_uncorrected_ < 0.005). Black outlines show language parcels. B. Mean selectivity within LH and RH language fROIs for language (blue) and cognitive load (yellow). C. Mean selectivity within LH and RH MD fROIs for cognitive load (yellow) and language (blue). Error bars denote standard error. Individual subject data points shown with hollow grey circles connected with grey lines. One-tailed t-test used to determine if selectivity is greater than zero; paired samples t-tests to compare selectivity across tasks. *p<0.05.

When we formally examined these same language fROIs for selectivity to the cognitive load during the working memory task using rmANOVA, we found a main effect of fROI in the LH (F(4,132)=5.52, p=3.86×10^-4^, η^2^=0.049), and a condition by fROI interaction (F(4,132)=3.25, p=0.014, η^2^=0.003), but no main effect of condition, F(1,33)=1.46, p=0.24, η^2^=0.003). Post-hoc t-tests revealed that the interaction effect was driven by the left orbital inferior frontal region, in which the response to the Easy condition was significantly higher than the Hard (t(33)=2.05, p=0.049, d=0.35, all other regions p>0.08, see **Table 2**), which is the opposite effect that we expect from the MD network (i.e. greater response to Hard, the more effortful condition, than Easy). There was no evidence of selectivity to cognitive load in the RH (main effect of fROI: F(3.26,107.61)=2.95, p=0.032, η^2^=0.028; but no significant effect of condition: F(1,33)=1.47, p=0.23, η^2^=0.0030; or interaction: F(3.23,106.75)=0.32, p=0.83, η^2^=0.00052).

**Table 2:**
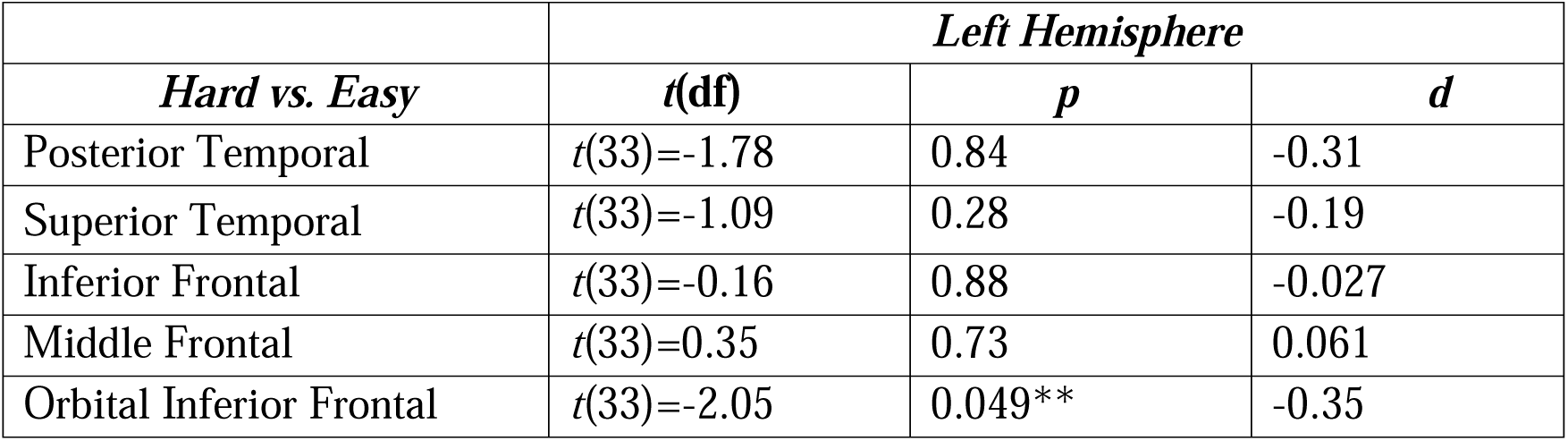
Specificity of language network fROIs. Note that only Left Hemisphere responses are tested because RH rmANOVA showed no effect of condition. Two-tailed, paired t-tests comparing responses to the Hard vs. Easy conditions (cognitive load selectivity) in the working memory task. T value, significance, and Cohen’s D effect sizes listed. *p<0.05, ** Bonferroni-Holm p<0.05, corrected for 5 comparisons.

|  | *Left Hemisphere* |  |  |
| --- | --- | --- | --- |
| *Hard vs. Easy* | *t(df)* | *p* | *d* |
| Posterior Temporal | $t(33)=-1.78$ | 0.84 | -0.31 |
| Superior Temporal | $t(33)=-1.09$ | 0.28 | -0.19 |
| Inferior Frontal | $t(33)=-0.16$ | 0.88 | -0.027 |
| Middle Frontal | $t(33)=0.35$ | 0.73 | 0.061 |
| Orbital Inferior Frontal | $t(33)=-2.05$ | 0.049** | -0.35 |

When directly comparing the selectivity of the language fROIs to language (Sn-Ns) and cognitive load (Hard-Easy), we find a significant main effect of task in both hemispheres (LH: task: F(1,30)=14.15, p=7.33×10^-4^, η^2^=0.15, nonsignificant fROI: F(4,120)=0.87, p=0.48, η^2^=0.005, significant interaction: F(4,120)=3.07, p=0.019, η^2^=0.015; RH: significant task: F(1,26)=5.00, p=0.034, η^2^=0.082, nonsignificant: fROI: F(4,124)=2.33, p=0.06, η^2^=0.010 and interaction F(3.21, 99.6)=1.92, p=0.13, η^2^=0.009; see **Figure 2B**), such that language selectivity is significantly higher than selectivity to cognitive load across fROIs (averaged LH fROIs: t(30)=3.76, p=7.33×10^-4^, d=0.68; averaged RH fROIs: t(31)=3.90, p=4.81×10^-4^, d=0.69). We see no evidence of selectivity to cognitive load in the language fROIs, and instead find evidence in support of specificity of the language network to the properties of high-level language. Note that for the analyses above we use all available datapoints with complete data for each analysis, but we also ran the same analyses on a subset of children motion-matched across the tasks and replicated the findings (dissociation between language and MD in language fROIs; see **Supplemental Results**).

To ensure that the language fROIs were not lacking cognitive load selectivity because this young sample of children had not yet developed this domain-general response, we also defined subject-specific fROIs for the MD network (see **Methods**) and assessed their selectivity to cognitive load. In a motion matched subset (two-tailed, paired samples t-test of framewise displacement in language and spatial working memory task; t(16)=-2.07, p=0.055) of subjects who completed two runs of the working memory task and performed well (>50% accuracy on hard trials; N=17), we examined the selectivity of the MD network (averaged across fROIs for each hemisphere) to cognitive load using a rmANCOVA of condition (Hard, Easy) with accuracy on the Hard trials as a covariate (see (Schettini et al., 2023)). We found a significant main effect of condition and an interaction between condition and accuracy in both hemispheres (LH: condition: F(1,15)=29.38, p=7.09×10^-5^, η^2^=0.10; interaction: F(1,15)=12.47, p=0.0030, η^2^=0.045; nonsignificant accuracy: F(1,15)=0.66, p=0.48, η^2^=0.40; RH: condition: F(1,15)=34.90, p=2.88×10^-4^, η^2^=0.12; interaction: F(1,15)=4.95, p=0.042, η^2^=0.019; nonsignificant accuracy: F(1,15)=2.06, p=0.17, η^2^=0.11), such that the response to Hard trials was significantly greater than Easy trials (LH: t(16)=4.14, p=7.75×10^-4^, d=1.00; RH: t(16)=5.29, p=7.31×10^-5^, d=1.28), suggesting that this sample has already developed the expected effortful response in the MD network. Further, the MD network did not show language selectivity in either hemisphere (i.e., no significant effect of condition: LH: F(2,30)=0.17, p=0.84, η^2^=0.004; RH: F(2,30)=0.29, p=0.75, η^2^=0.006; results are unchanged when including working memory task accuracy as covariate). As expected, the MD network is significantly more selective to cognitive load than language (ANOVA of selectivity by task, LH: F(1,16)=5.65, p=0.03, η^2^=19; RH: F(1,16)=8.37, p=0.011, η^2^=0.25; follow-up two-tailed, paired samples t-tests of selectivity to cognitive load vs language: LH: t(16)=2.38, p=0.03; RH: t(16)=2.89, p=0.011; see **Figure 2C**). Combined with the language network analysis above, these results suggest that language is functionally dissociated from domain-general processes that recruit the MD network.

Finally, if there was a progressive change between the language network’s sensitivity to language and cognitive load across development (see example subjects’ activation at two timepoints **Figure 3A**), we might expect the language network’s responses to the linguistic and the working memory task to be related. However, there was not a systematic shift in which language activation increased as cognitive load responses decreased; instead language and selectivity to cognitive load in the language fROIs were not correlated (LH: r(34)=-0.081, p=0.64; RH: r(34)=0.034, p=0.85) in the cross-sectional sample. These findings did not change when controlling for age (partial correlations; LH: r(33)=0.012, p=0.95; RH: r(33)=0.044, p=0.80, **Figure 3B**). Examining the longitudinal changes to these responses within individual, we also did not find a relationship between changes in language and selectivity to cognitive load within an individual in the language fROIs (LH: r(15)=-0.14, p=0.59; RH: r(15)=-0.098; p=0.71; partial correlation controlling for age at timepoint one: LH: r(14)=0.32, p=0.23; RH: r(14)=0.11, p=0.67, **Figure 3C**). These results suggests that changes in the language fROIs selectivity to language across ages and within an individual are not related to either a systematic increase or decrease in these regions’ responsiveness to a domain-general task.

**Figure 3:**
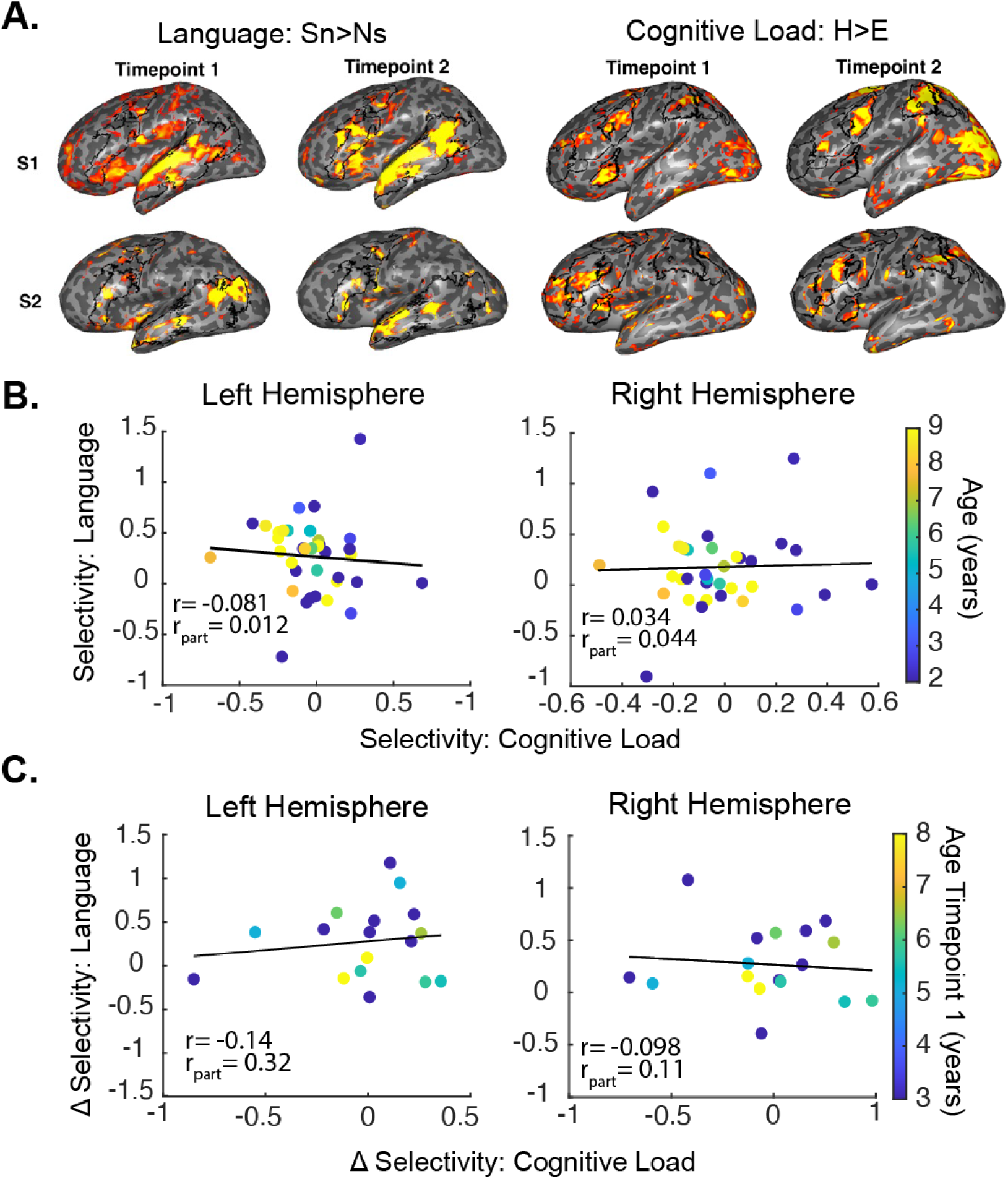
A. Language (left) and cognitive load (right) activation for two example subjects at timepoint one and two (subject 1 age: TP1=5.85, TP2: 7.79 years; subject 2 age: TP1: 4.56, TP2: 7.43 years; threshold at P_uncorrected_ < 0.005). Black outlines show language parcels (left) and MD parcels (right). B. Scatter plot with regression line for selectivity to language and cognitive load in the LH and RH language fROIs (N=36). C. Scatter plot with regression line for selectivity to language and cognitive load in the longitudinal sample (N=17) in the LH and RH language fROIs. Individual subject data points colored by age (yellow is oldest, and blue is youngest). R values for correlation and partial correlations (controlling for age) are listed in each scatter plot; no correlations were significant.

### Is the high-level language network differentiated from domain-general cortex in connectivity?

Seeing that the language and MD networks are dissociable in their functional specificity, we next examined if these networks are dissociable in their resting-state functional connectivity. Both networks show characteristically significant bilateral within-network connectivity (two-tailed, one-sample *t*-tests, **Table 3**, **Figure 4**), with higher within-network connectivity in the LH compared to the RH for the language network (paired samples *t*-test: *t*(31)=2.65, p=0.012, d=0.47) and no hemisphere difference for the MD network (*t*(31)=-0.11, p=0.91, d=-0.019). Perhaps unexpectedly, given the differentiation of language and MD in function, we did observe significant between-network (language to MD) connectivity (**Table 3)**, although this between-network connectivity was significantly lower than the within-network connectivity of both networks (within vs between: Language: LH: *t*(31)=5.10, p=1.58 x 10^-5^, d=0.90; RH: *t*(31)=4.55, p=7.72 x 10^-5^, d=0.80; MD: LH: *t*(31)=3.73, p=7.66×10^-4^, d=0.66; RH: *t*(31)=3.56, p=0.0012, d=0.63).

**Figure 4:**
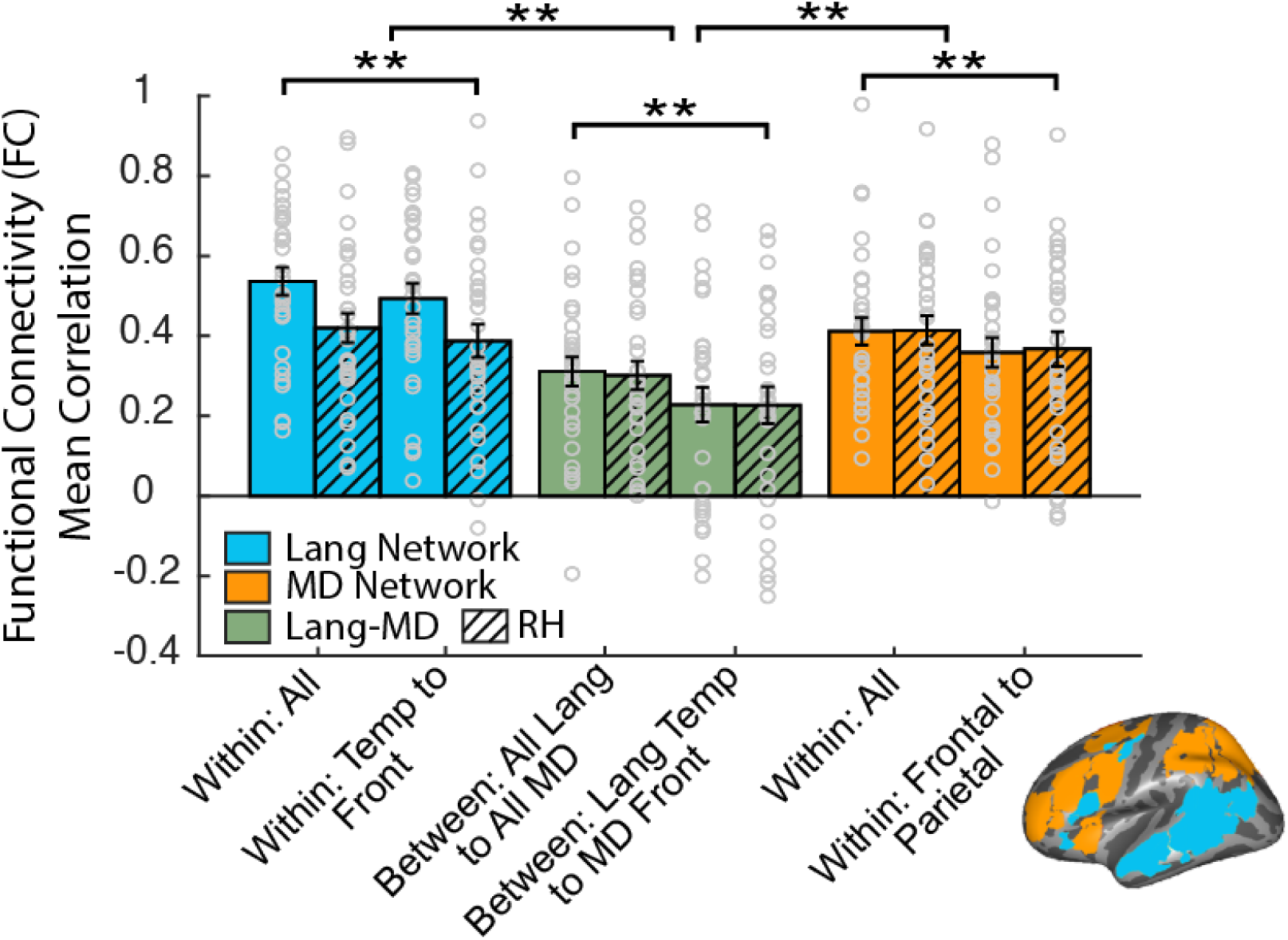
Within- and between-network functional connectivity (FC) of the language and MD fROIs. One-sample, two-tailed t-tests tested if FC was significantly greater than zero. Repeated-measures t-tests were used to compare FC across within- and between-network connectivity. Grouped bars indicate all under the group are significant. Group comparisons indicate that within bars are significantly higher than between bars (e.g., Within all LH for language significantly higher than between all language to and MD). Asterisks indicate level of significance (* p < 0.05; ** Bonferroni-Holm p < .05). Error bars denote standard error. Individual subject data points shown with hollow grey circles. Inset shows parcel search spaces used for creating fROIs for the language (blue) and MD network (orange).

**Table 3:** Within- and between-network connectivity of the language and MD network in N=32 individuals. Two-tailed, one-sample t-tests compare connectivity to zero. T value, significance, and Cohen’s D effect sizes listed. *p<0.05, ** Bonferroni-Holm p<0.05, corrected for six comparisons.

|  | Left Hemisphere |  |  | Right Hemisphere |  |  |
| --- | --- | --- | --- | --- | --- | --- |
| Within-Network | *t* | *p* | *d* | *t* | *p* | *d* |
| Language | 15.50 | $3.75 \times 10^{-16}^{**}$ | 2.74 | 11.38 | $1.353 \times 10^{-12}^{**}$ | 2.01 |
| Lang: Frontal to Temporal | 13.06 | $3.83 \times 10^{-14}^{**}$ | 2.31 | 9.33 | $1.64 \times 10^{-10}^{**}$ | 1.65 |
| Multiple Demand | 12.15 | $2.50 \times 10^{-13}^{**}$ | 2.15 | 11.37 | $1.38 \times 10^{-12}^{**}$ | 2.01 |
| MD: Frontal to Parietal | 9.74 | $5.85 \times 10^{-11}^{**}$ | 1.72 | 8.48 | $1.41 \times 10^{-9}^{**}$ | 1.50 |
| **Between-Network** | ***t*** | ***p*** | ***d*** | ***t*** | ***p*** | ***d*** |
| Language-MD | 8.48 | $1.49 \times 10^{-9}^{**}$ | 1.65 | 8.38 | $1.83 \times 10^{-9}^{**}$ | 1.48 |
| Lang. Temporal to MD Frontal | 5.27 | $9.79 \times 10^{-6}^{**}$ | 0.93 | 4.95 | $2.47 \times 10^{-5}^{**}$ | 0.88 |

Next, we examined age-related changes in connectivity. The within-network LH language connectivity was positively related to age in this cross-sectional sample (All LH: r(30)=0.42, p=0.018; LH Temporal to Frontal: r(30)=0.42, p=0.018, both n.s. after correcting for multiple comparisons; All RH: r=0.12, p=0.50; RH Temporal to Frontal: r(30)=0.073, p=0.69). The within-network MD connectivity was also positively related to age (All LH: r(30)=0.41, p=0.021; LH Frontal to Parietal: r(30)=0.42, p=0.029; All RH: r(30)=0.39, p=0.029; RH Frontal to Parietal: r(30)=0.40, p=0.025; all but RH n.s. after correction for multiple comparisons). The between-network connectivity was not significantly associated with age (LH: r(30)=0.17; p=0.37; RH: r(30)=0.10, p=0.59). Importantly, there was also no relationship between age and the difference in within- vs. between-network connectivity of language and MD regions (within language - between: LH: r(30)=0.19, p=0.30, RH: r(30)=0.038, p=0.84; within MD - between: LH: r(30)=0.29, p=0.11; RH: r(30)=0.33, p=0.06), suggesting that age-related increases in connectivity of the language network were not related to ongoing dissociation from the MD network; and vice-versa (i.e., that age-related increases in the MD network are not related to dissociation from the language network). Finally, there were no age-related changes in between-network connectivity.

We also examined changes in functional connectivity across time longitudinally within subjects (two-tailed, paired-samples t-test, timepoint one compared with timepoint two). We did not see any changes in within-network functional connectivity for either the language (All LH: *t*(13)=0.76, p=0.46, d=0.20; LH Temporal to Frontal: *t*(13)=0.36, p=0.72, d=0.097; All RH: *t*(13)=0.31, p=0.76, d=0.083; RH Temporal to Frontal: *t*(13)=0.24, p=0.82, d=0.64) or MD network (All LH: *t*(13)=-0.74, p=0.47, d=-0.20; LH Frontal to Parietal: *t*(13)=-0.44, p=0.66, d=-0.29; All RH: *t*(13)=-1.07, p=0.30, d=-0.29; RH Frontal to Parietal: *t*(13)=-1.14, p=0.27, d=-0.29). We did not observe changes in between-network connectivity of language and MD over time within subject (LH: *t(*13)=-0.81, p=0.43, d=-0.22; LH Language Temporal to MD Frontal: *t*(13)=-0.64, p=0.53, d=-0.17; RH: *t*(13)=-1.34, p=0.20, d=-0.36; RH Language Temporal to MD Frontal: *t*(13)=-0.63, p=0.54, d=-0.17). Thus, we see alignment of our cross-sectional and longitudinal results, with no robust age-related changes in language network connectivity, MD network connectivity, or between-network connectivity of language and MD.

## Discussion

Understanding the development of the language network and its specificity in function and connectivity will help us better understand the neural mechanisms that underlie uniquely human cognition. In this paper, we examined whether the language network shows early specialization for language, and if there is evidence of language cortex emerging from domain-general cortex. We used subject-specific fROIs to account for individual variability in the precise location of the language network across individuals and collected multiple runs of data, matching the methodological rigor of fMRI work in adults. We also removed the confound of task difficulty by using a passive language task (widely validated in adults to activate the language network, regardless of language; Malik-Moraleda et al., 2021), that contrasts a meaningful Sentences condition with a Nonsense sentences condition. This Nonsense condition importantly removes two key aspects of language – semantic meanings and syntatic relationships – while keeping intact prosody and rythym of speech by speaking meaningless words as if they were sentences. And, for the first time to the best of our knowledge, we compare activation to a linguistic vs. non-linguistic task to characterize the development and dissociation of the language network in a young sample of children.

We find that the language network is selective to language in both frontal and temporal language regions and left-lateralized, in line with some previous work in older children (e.g. Charbonnier et al., 2020; Enge et al., 2020; Olulade et al., 2020; Weiss-Croft & Baldeweg, 2015). However, here we directly compare linguistic responses in each child to their neural responses while performing a nonverbal yet demanding task to see if the language network is domain-specific or whether it emerges from domain-general processing used for other computations (it does not). And while selectivity to high-level linguistic properties (e.g. semantics and syntax) increase with age and across time within our longitudinal subjects, we do not find evidence of a progressive shift from domain-general to domain-specific functioning in this sample of young children. Finally, by additionally examining resting-state data in both our cross-sectional and longitudinal samples, we found that the language and MD networks are also dissociable in their connectivity over time, with both the language and MD networks showing more within-network than between network connectivity, and stable connectivity across ages and longitudinally within subjects.

Using two separate tasks and subject-specific fROIs, we find a double-dissociation between language and MD networks in a young sample of children. These results are contrary to hypotheses that language development is part of either general skill-learning (as has been suggested by (Chater & Christiansen, 2018; McCauley, 2020; Saffran & Thiessen, 2007)) or is bootstrapped by executive control (Edgin et al., 2015). For example, combinatorial processing is one of the hallmarks of language but possibly used for other high-level computations as well. Instead, the language network is not responsive to a spatial working memory task that localizes the MD network, mirroring prior work in adults (Fedorenko et al., 2012) and suggesting that language cortex is indeed domain-specific for linguistic content even in young children.

Importantly, even though we did not see any evidence of load-based domain-general responses in the language network, it is possible that there is a *progessive shift* across time, in which case the language network would begin as sensitive to domain-general processing, which may act as a scaffolding mechanisms to support language specificity through time and greater language exposure (see neuroconstructivism hypothesis, (Karmiloff-Smith, 2015)). In this case, we would expect to see changes in sensitivity to cognitive load during the working memory task longitudinally within subject, such that there is an initial response to both cognitive load load and language at young ages (reflecting domain-general processing), followed by a decrease in load-based responses and increase in language selectivity (reflecting increase in domain-specific processing). This is not the case in the cross-sectional subjects or longitudinally within children. We see a significant increase in language selectivity in the left hemisphere with age cross-sectionally and longitudinally (aligning with prior work on receptive language; Berl et al., 2012, Szaflarski et al., 2006). Howevever, we do not see an initial sensitivity to working memory load within the language network in our young subjects; nor we do increases in this load-based response in our longitudinal sample or any correlation of these responses. We further find that the MD network (as defined using the working memory task) shows the expected load-based preference but not language preferences (see also Schettini et al., 2023). Overall, this evidence suggests that the language and MD networks develop independently in young children.

Prior work suggests that language responses are more bilateral in children and become increasingly LH lateralized with age, presumably through inter-hemispheric connections that inhibit RH (or non-dominant hemisphere) responses over time (e.g. (Friederici et al., 2011; Holland et al., 2007)). In contrast to this theory, we do not see decreasing language selectivity in the RH language regions in young children. In more recent prior work, Olulade and colleagues (Olulade et al., 2020) using cross-sectional analysis in slightly older ages (4-10 years, N=24) found decreasing language selectivity in the RH inferior frontal gyrus (IFG) with age and the oldest child group showed no such activation on the right. It is possible that they could not rule out confounds of spatial precision (i.e., they do not define the language network for each individual) or effort-based attention (i.e., no additional task and varying response difficulty to conditions within their language task). Importantly, our probabilistic atlas shows this sample of young children have very low overlap in RH language activation, making it even more important to define language regions within subjects. Additionally, because the IFG houses both language and MD regions, their observed decreases in “language” activation in the right IFG may actually reflect decreasing responses within language regions to effort-based attention or decreasing responses within MD regions to linguistic content or any other combination of possibilities. Our use of subject-specific fROIs and our differentiation of language and domain-general cortex using two tasks in both cross-sectional and longitudinal samples allowed us to better characterize the development of the domain-specific language network in young children, showing that if the language network starts out more bilaterally, then it would be earlier than the youngest ages we study here (i.e. less than 2-3 years of age). Further, the present results are more in line with adult literature showing robust RH language activation, just with more dominant activation in the LH (Lipkin et al., 2022).

While no previous study empirically investigated the neural dissociation of linguistic vs. non-linguistic content in children, one review has examined the relationship between cognitive flexibility, inner speech, and language in children and young adults, finding that while inner-speech and language are not required for cognitive flexibility, it can enhance performance on a variety of cognitive flexibility tasks. We do not rule out this possibility and suggest that perhaps this interaction between cognitive flexibility and language is supported by the significant connectivity between the language and MD network. However, this connectivity was dwarfed by within-network language connectivity, suggesting that the language network is indeed largely independent of other adjacent cognitive networks. Additionally, our within-network language connectivity findings mimic what previous studies show in adults, with language typically left-lateralized and with significantly higher left than right within-network FC. Finally, when examining the connectivity within and between the language and MD networks, we found no reliable increases in within- vs. between-network FC cross-sectionally and longitudinally. This suggests that specificity of connections may be more static than neural specialization, at least for the domain of language, supporting previous work testing the connectivity-fingerprints hypothesis, where connectivity precedes and perhaps drives the functional organization and specificity of cortex (Hannagan et al., 2015; Li et al., 2020; Saygin et al., 2016; Yu et al., 2020). Prior longitudinal work has suggested that changes in within-network connectivity is associated with changes in language performance, as opposed to age (Xiao, Brauer, et al., 2016; Xiao, Friederici, et al., 2016); however, we could not test that hypothesis with our data.

Multiple questions remain. First, we do not examine behavioral metrics of language development, and are therefore unable to differentiate between changes in language selectivity that are due to maturation vs. language skill development. Additionally, using a passive language task does not allow us to understand development of specific language abilties (grammar, comprehension, or vocabulary). While basic language skills are largely developed in our youngest cohort, it would be interesting to examine the continued development of this network in parallel to gains in particular linguistic skills (e.g. (Abrahams et al., 2003; Ardila, 2011, 2012). Further, this work is limited by a lack of an *a priori* power analysis. However our overall sample size is on par with previous developmental cognitive neuroscience studies. Our core sample includes 45 children with two runs of the language task, 36 with at least one run of the spatial working memory task, 19 longitudinal subjects with both tasks, which is comparable or higher than prior work (e.g., (Olulade et al., 2020) N=24 under age 10; only one of the 27 studies included in the (Enge et al., 2020) meta-analysis on language comprehension in children had a larger core sample: N=57 (Berl et al., 2014) but their mean age (8.9) which is on par with our maximum age (9.1)). Despite this, language development should continue to be examined as more data is collected, and in even younger samples. Finally, it would be of interest for future work to continue to examine the relationship between lateralization, selectivity, and connectivity of the language network, preferably in younger longitudinal samples.

## Methods

### Participants

As part of an ongoing project exploring the relationship between functional organization of the human brain and connectivity, participants completed a battery of functional tasks in the scanner. Data were collected from 47 children. Of the 47 children, 45 children completed structural imaging and two runs of the language localizer task during at least one timepoint (23 female, 32 right-handed, mean (sd) age=5.92 (1.70) years, age min and max = 36.5 months-9.07 years, 35 White, 3 Bi-Racial, 2 Asian, 4 Black, 1 Latino) and were used in the cross-sectional analysis. A subset of this group completed at least one run of the spatial working memory task (used to localize the MD network, N=35, mean (sd) age=6.26 (1.64) years). Of the 45 children, 32 completed resting state scans and at least one run of both the language and working memory task, so were therefore included in the resting state analysis.

33 children returned for at least one subsequent, yearly timepoint and 23 of those children (12 female, 19 right handed, 18 White, 2 Bi-Racial, 1 Black, 1 Asian, 1 Latino, timepoint 1 (TP1): mean (sd) age=5.50 (1.28) years, age min and max = 3.95-7.93 years; timepoint 2 (TP2): mean (sd) age =7.00 (1.41) years, age min and max = 4.74-9.07 years, mean time between appointments=1.50 years) completed two runs of the language localizer at their latest timepoint, at least one run of the language localizer at an earlier timepoint, and passed data quality checks and were therefore included in the longitudinal analysis. 19 of these children completed at least one run of the MD task at each timepoint, as well as a resting state and structural imaging and were included in the MD longitudinal analysis. For children who completed all scans at multiple timepoints, the earliest timepoint was prioritized for cross-sectional analysis.

This study was approved by the institutional review boards at The Ohio State University. Parents gave written consent and children gave verbal assent to participate.

### Data acquisition and Preprocessing

To acquire data, children arrived approximately one hour before the scanning appointment to practice the ‘games’ (tasks), to practice lying still in a mock scanner, and to become comfortable with the scanning environment. Children had the option to bring a stuffed animal to the scanning session, to keep them company during the scan. Additionally, one experimenter remained in the scanning room with the child throughout the procedure to ensure the child understood the rules of each game, was lying still, and paid attention to each task. In the case that the child was confused, not responding to the game, or needed a break, the experimenter in the room would alert study staff, the scan would be paused until the child was ready to resume. These procedures helped ensure children were still during the scan, felt comfortable, and willing to return for additional longitudinal appointments.

Anatomical: Images were acquired on a Siemens Prisma 3T scanner with a 32-channel phase array receiver head coil. Foam padding was used for head stabilization and increased comfort throughout all scanning. A whole-head, high-resolution T1-weighted magnetization-prepared rapid acquisition with gradient echo (MPRAGE) scan was acquired (repetition time (TR) = 2300 ms, echo time (TE) = 2.9ms, voxel resolution = 1.00 mm^3^). Data were analyzed with Freesurfer v.6.0.0, FsFast, FSL, and custom Matlab (version R2020a) code. A semi-automated processing stream (recon-all from Freesurfer) was used for structural MRI data processing. Major preprocessing steps include intensity correction, skull strip, surface co-registration, spatial smoothing, white matter and subcortical segmentation, and cortical parcellation. The cortical surface of each subject was examined following segmentation and was manually adjusted when necessary. Cortical gray matter masks were created in native anatomy for each subject.

Functional: Images for the localizer tasks (see below *Functional Localizer* section), as well as resting-state scan, were acquired with multiband 4x accelerated the echo-planar imaging (EPI) sequence: TR=1000ms, TE=28ms, voxel resolution = 2×2×3 mm. For localizer tasks, data was preprocessed with motion correction, smoothing (FWHM =4mm), and bbregister (Greve & Fischl, 2009) was used to register functional data to the subject’s anatomical space. All data were analyzed in each individual’s native anatomy. For resting-state scan, Freesurfer’s FS-Fast preprocessing (https://surfer.nmr.mgh.harvard.edu/fswiki/FsFastAnlysisBySteps) was used to complete motion correction and smoothing (FWHM=3mm). Additional preprocessing included linear interpolation for spikes over 0.5mm and a bandpass filter at 0.009-0.08 Hz. All functional connectivity (FC) analyses were performed in native functional space. During the resting state scan, children were instructed to “have a staring contest” with a white crosshair on a black background. The experimenter in the scan room watched to see if the child was awake/following instructions. The resting-state scan lasted approximately 5 minutes and consisted of 290 volumes for a single run.

### Functional Localizer

Language Localizer: A language localizer task (Fedorenko et al., 2010) was used to functionally locate the regions of the brain responsive to the lexical and structural properties of language. Participants listened to blocks of meaningful Sentences, Nonsense sentences (constructed from phonemically intact nonsense words that control for features for speech, such as phonology and prosody), and Texturized speech (controlling for low-level auditory features). Each run consisted of 4 blocks of each condition and three 14-second fixation blocks. Each block contained three trials (6 seconds each); each trial ended with a visual queue to press a button. Each participant finished 2 runs of the task whenever possible.

Multiple-demand localizer: A spatial working memory task (Fedorenko et al., 2013) with blocked Easy and Hard conditions (task difficulty adjusted by age; low, medium and high load versions of the task) were used to functionally define brain regions associated with the domain-general MD network and examine the brain’s response to cognitive load. In each trial, participants viewed a grid of 9 (low load) or 12 (medium and high load) boxes. For the Easy condition of the medium load, two grid patterns are shown sequentially, with a different box highlighted in blue at a specific location of the grid and the child must remember the location of these blue boxes (Hard blocks have more blue boxes than Easy blocks). Then two grids appear side-by-side, and the child indicates via a button press (using their left or right hand) if the pattern of blue boxes in the left or right grid matches the previously presented grids. Task difficulty was adjusted by age (low [4-5 years], medium [6-7 years], and high [8+ years] load), and the low load version of the task had a longer response duration, to allow more time for children to answer. Children would practice their age defined level before the scan session and the level would be adjusted (made easier if child missed most questions, or more difficult if the child showed near-perfect performance) if necessary.

### Statistical Analysis

Task fMRI Analysis: Preprocessed data were entered into a volume-based, first-level general linear model (GLM) analysis. A regressor was entered for each condition of interest (language localizer: meaningful sentences (Sn), nonsense sentences (Ns), texturized speech (Tx); MD localizer: Hard and Easy conditions), and six motion parameters were included as nuisance regressors. Timepoints with motion exceeded 1mm framewise displacement (total motion between two consecutive timepoints) were included as an additional nuisance regressor. A block design with a standard boxcar function convolved with the canonical hemodynamic response function (standard gamma function (d=2.25 and t=1.25)) was used. Relevant contrasts (language localizer: Sn>Ns, Sn>Tx, and Ns>Tx; spatial working memory localizer: Hard > Easy) were included in the level 1 processing. The GLM was completed in 2×2×2 mm^3^ native anatomical space and then was resampled to 1×1×1mm^3^. The resulting beta estimates and contrast maps were used for further analyses. Across both localizers, any run that exceeded the threshold of more than 25% of timepoints with greater than 1mm total vector motion between subsequent timepoints were excluded from the analysis.

### Quantifying Motion

For each task and each subject, the motion estimates (x, y, z, roll, pitch, yaw) for each subject were used to calculate the framewise displacement at each timepoint, excluding timepoints with greater than 1mm motion, which were included as nuisance regressors in first-level GLM analysis. The mean framewise displacement for each subject was calculated for each run of the language and spatial working memory task, then averaged across runs to create one framewise displacement number per subject per task. Pairwise t-tests were used to compare mean framewise displacement across tasks. Motion was also compared across timepoints for each task for the longitudinal subjects. Mean framewise displacement was also included as a covariate in correlations between selectivity (or percent signal change) and age in the cross-sectional sample (see below).

### Generating a Probabilistic Atlas

To quantify the number of children in our sample that show language activation, a probabilistic atlas was generated from the Sentences > Nonsense contrast. The second-level GLM significance maps (combined from run 1 and run 2) of the language task was registered from individual subject volume anatomical space to the surface (using Freesurfer’s mri_vol2surf function) and then to fs-average space (using Freesurfer’s mri_surf2surf function, https://surfer.nmr.mgh.harvard.edu/fswiki/FsAverage). Each registered significance map was thresholded at p < 0.01, binarized, and summed to show the number of subjects with significant activation at each voxel.

### Defining subject-specific functional regions of interest

Ten frontotemporal language network (five in each hemisphere; posterior temporal, superior temporal, inferior frontal, middle frontal, and orbital inferior frontal) subject-specific fROIs were created using the group-constrained subject-specific approach (Fedorenko et al., 2010). These parcels (Fedorenko et al., 2010; https://evlab.mit.edu/funcloc/), which contain the regions typically activated by high-level language based on prior adult studies, were used as spatially constrained search spaces for defining individual subject fROIs based on the Sentences > Nonsense speech contrast. Upon visual inspection, the frontal parcels did not fully capture some children’s hotspots of frontal language activation (see **Supplemental Methods**). Therefore, the frontal parcels were dilated by 3 (with Freesurfer’s mri_binarize command), which ensured hotspots of activation across subjects were covered and helped make the frontal parcels more comparable in size to the temporal parcels. These parcels were all registered to each participant’s native anatomy using CVS registration (Zöllei et al., 2010). Participant-specific fMRI contrasts maps from one run of the GLM analysis were masked to only include gray matter voxels, and the intersection of these contrast maps with the parcels were used to define the fROIs for each individual subject. We defined fROIs as the top 10% of voxels that fell within each parcel/search space (for language fROIs we used contrast of Sentences > Nonsense). The beta estimates from an independent run of the language localizer task were then used to extract the percent signal change (PSC) for each condition (Sentences, Nonsense, Texturized) within the subject-specific fROIs. Each subject completed two runs of the language task in a single timepoint; therefore, one run was used to define fROIs and the other run was used to extract the PSC, then this was repeated using opposite run orders and PSC results were averaged together for each participant. Any PSC values that exceed +/- 3 standard deviation across subjects within an fROI for each condition were marked as outliers and removed from subsequent analyses. This was done separately for cross-sectional, motion matched, and longitudinal samples. Using the same fROIs defined from the independent language task, we extracted PSCs from the spatial working memory task for the Hard and Easy conditions (averaged across runs for children who completed more than one run), to compare responses within the language network between the two tasks. Additionally, we used the same method to define MD network fROIs (10 bilateral, Fedorenko et al., 2013) and extracted PSC from an independent run of the spatial working memory task to examine the MD network selectivity to the domain-general Hard > Easy cognitive load effect and from at least one run of the language task.

For the 24 longitudinal subjects, language fROIs were created for the later timepoint following the same procedure (for Sentences > Nonsense, 20% threshold) as the cross-sectional data. Then, the overlapping voxels between these two fROIs at the later timepoint (20% threshold used to increase likelihood of overlap across fROIs) were identified and selected to form the new fROI that was registered to each subject’s earlier timepoint anatomical space using Freesurfer’s CVS registration (Zöllei et al., 2010), and PSCs to each condition of interest was extracted from the earlier timepoint’s beta estimates from the first-level GLM. Most subjects had overlap in their TP2 fROIs; however, subjects with less than 10% of the maximum overlap of all subjects were excluded, to ensure a robust and reliable fROI for each individual at the earliest timepoint. Finally, the registration between timepoints was visually checked for all subjects. One subject had poor registration between the two timepoints (leading to the fROIs mapped from the latest to earliest timepoint falling outside the brain), and was therefore excluded, leaving 23 subjects with longitudinal data for further analysis. For the final analysis, all fROIs had at least 19 subjects with enough overlap to create a TP1 fROI (all subjects with fROI: LH posterior temporal, LH superior temporal, LH inferior frontal, RH inferior frontal; 1 subject missing fROI: LH orbital inferior frontal, RH posterior temporal, RH superior temporal, RH middle frontal; 2 subjects missing: RH orbital inferior frontal; 4 subjects missing: LH: middle frontal).

### Selectivity

To quantify an individual’s selectivity to language or MD within each fROI, we calculated selectivity by adding the most negative PSC across conditions within a task (to force the lowest value to zero to “correct for potential bias induced by negative activation” (Simmons et al., 2007; Szwed et al., 2011) to all conditions by subject and subtracting the PSC of the control condition of interest from the condition of interest. Selectivity reflects the difference between the condition of interest and the control condition for each task.

1. Language Selectivity = *PSC_sentences_* - *PSC_Nonsense_*
2. Multiple Demand Selectivity = *PSC_Hard_* - *PSC_Easy_*

### Determining Selectivity and Specificity of fROIs

A series of repeated-measures analysis of variance (rmANOVA) and repeated-measures analysis of covariance (rmANCOVA) were used to determine selectivity specificity of language-network fROIs to conditions of interest across both tasks, and any changes with age. All rmANOVA were implemented in RStudio (version 1.4.1717, R version 4.1.1), using the anova_test function (rstatix, lme4 and lmeTest packages). In any cases that the Mauchly’s Test of Sphericity showed a significant effect indicating that sphericity was violated, a Greenhouse-Geisser epsilon correction was applied, and all reported effects remain significant following this correction and corrected p-values are reported. Generally, following significant interaction, post-hoc, two-tailed, paired samples t-tests were conducted with Bonferroni-Holm (Holm, 1979) multiple comparison corrected by the number of fROIs within hemisphere. All rmANOVAs were run separately for each hemisphere, and any subject with missing data (due to outlier exclusions) in any fROI for that hemisphere was excluded for that analysis.

To determine the selectivity of the language network to the properties of high-level language we used a rmANOVA of fROI (for five fROIs in the hemisphere) by condition to explain PSC during the language task. A significant effect of condition in which the Sn response is higher than both the Ns and Tx would indicate a selectivity of the language fROIs to the high-level properties of language. Additionally, the laterality of the language network was assessed using a rmANOVA of fROI by hemisphere explaining language selectivity. A significant main effect of hemisphere, in which the selectivity of the LH fROIs is significantly higher than the RH fROIs (as determined with paired samples t-tests) would indicate left-laterality in this sample.

To understand any age-related changes in the selectivity and specificity of the language network, we used a condition by fROI rmANCOVA with age as a covariate, separate for each hemisphere. In the case of a significant interaction between age and condition, post-hoc Pearson correlations were used to examine changes in each condition (averaged across ROIs within a hemisphere) with age, and follow-up partial correlations controlling for mean framewise displacement were also run. We also examined age-related changes in the longitudinal sample with a condition by timepoint rmANOVA (by hemisphere). Post-hoc t-tests were used to determine significant difference over time for each condition following significant timepoint by condition interactions.

To determine the specificity of the language network to language, we ran a similar rmANOVA of fROI by condition, this time with the PSC to conditions in the spatial working memory task as the outcome variable. A significant effect of condition, in which the PSC to the Hard condition was significantly higher than the Easy, would indicate a selectivity of the language fROIs to the working memory; however, a nonsignificant main effect of condition or condition effect driven by a higher Easy than Hard response would indicate that the language network is not responsive to this working memory task. We additionally examined the specificity of the language network by comparing the language and working memory selectivity indices within the language network using a rmANOVA of fROI by task; in which a significant main effect of task in which language selectivity is significantly higher than MD would support the idea that the language network is both sensitive and specific for language in this young sample; whereas a nonsignificant task effect would suggest language and working memory responses are intertwined in this sample.

We repeated the above analyses for the MD fROIs as well, to examine the responsiveness to the MD network (one average value across fROIs for each condition and each hemisphere) to both tasks. One rmANOVA (for each hemisphere) was used to examine differences in PSC across conditions (Hard and Easy) in the working memory task. A separate rmANOVA was run to examine PSC during the language task. Finally, a third rmANOVA was used to compare selectivity of the MD fROIs to both language and working memory. Similar to when examining the language fROIs, if the MD fROIs show selectivity to working memory (significant condition effect for Hard and Easy condition), specificity (no significant condition effect to the language conditions), and a task effect with greater working memory than language selectivity, that would indicate the MD fROIs are responsible for domain general processes like working memory, but not language.

Finally, we examined whether there was a relationship between changes in the language network’s sensitivity to language and working memory across subjects. We used a correlation of language selectivity and working memory selectivity to assess this and additionally controlled for age in a partial correlation. We also examined this within-subject by correlating change in language selectivity with change in working memory selectivity, and we completed a follow-up partial correlation controlling for age at timepoint one. If language and working memory selectivity were correlated, this would suggest language and working memory selectivity were related across development (perhaps progressively shifting, in which an initial response to the Hard>Easy is replaced by language specific responses at older ages or across time); whereas no relationship between the two would suggest independence of the two networks.

### Within- and Between-network Connectivity

For 32 unique children, the mean time course was extracted from five bilateral language fROIs (Fedorenko et al., 2010) (as defined above with the Sentences > Nonsense 10% threshold) and 10 bilateral frontoparietal MD fROIs (as defined above with the spatial working memory Hard > Easy 10% threshold; (Fedorenko et al., 2013) registered to the subject-specific resting state data. Functional connectivity (FC) was calculated using Pearson’s correlations between each pair of fROIs. To generate normally distributed values, each FC value was Fisher z-transformed. We examined within and between-network connectivity of the language and MD fROIs, as well as comparing the within vs. between connectivity of the networks. Within-network connectivity was tested by comparing the mean connectivity (by hemisphere and between lobes for temporal to frontal (language) and temporal to parietal (MD network) value with zero using two-tailed, independent samples *t*-tests. We also compared connectivity by hemisphere to examine laterality (two-tailed paired-samples *t*-test for each network). Between network connectivity (by hemisphere and for temporal language to MD frontal) was also tested using two-tailed, independent samples *t*-tests. Within-network connectivity of both language and MD were compared with between-network connectivity using two-tailed, paired *t*-tests (grouped by hemisphere and by lobe). Bonferroni-holm multiple comparison correction was applied for all within-network and between-network tests. Pearson’s correlations were used to assess the relationship between within-network FC (by hemisphere) and age, between-network FC (by hemisphere) and age, and the difference of within vs between network connectivity and age for both the language and MD network (by hemisphere for each).

For the 17 longitudinal subjects with two runs of both tasks at the later timepoint as well as resting state data, longitudinal fROIs were used (see above) to examine changes in functional connectivity. Registration between timepoints was visually examined for each subject, and three subjects were removed because of poor registration to resting state data. The mean time course was extracted, and FC was calculated. Changes in FC were assessed using two-tailed, paired samples t-tests comparing the FC in timepoint one with timepoint two for within network language and MD, as well as between network FC.

### Data and Code Availability

Please contact Dr. Zeynep Saygin for data, code, and software requests. MATLAB, Bash, and FreeSurfer software were used to process these data. De-identified data used for figures and tables, as well as all processing code will be available on GitHub upon publication (https://github.com/SayginLab). To access raw study data for research purposes, researchers can sign a data use agreement between institutions and project proposal as required by The Ohio State University. Commercial use of the raw study data was not included as part of the current IRB-approved project and is therefore restricted to research use.

## Acknowledgements

We are grateful to the families who generously supported this work. We would like to thank members of the Saygin Developmental Cognitive Neuroscience Lab for data collection. We would like to acknowledge the support from Center for Cognitive and Behavioral Brain Imaging (CCBBI) and Ohio Supercomputer Center (OSC). This material is based upon work supported by the National Science Foundation Graduate Research Fellowship (awarded to K.J.H.) under Grant No. DGE-1343012. Z.M.S. was supported by the Alfred P. Sloan Foundation, OSU’s College of Arts & Sciences, and the Chronic Brain Injury initiative at OSU.

## References

Abrahams, S., Goldstein, L. H., Simmons, A., Brammer, M. J., Williams, S. C. R., Giampietro, V. P., Andrew, C. M., & Leigh, P. N. (2003). Functional magnetic resonance imaging of verbal fluency and confrontation naming using compressed image acquisition to permit overt responses. Human Brain Mapping, 20(1), 29–40. 10.1002/hbm.10126

Amunts, K., Schleicher, A., Bürgel, U., Mohlberg, H., Uylings, H. B., & Zilles, K. (1999). Broca’s region revisited: Cytoarchitecture and intersubject variability. The Journal of Comparative Neurology, 412(2), 319–341. 10.1002/(sici)1096-9861(19990920)412:2<319::aid-cne10>3.0.co;2-7

Ardila, A. (2011). There are Two Different Language Systems in the Brain. Journal of Behavioral and Brain Science, 01(02), Article 02. 10.4236/jbbs.2011.12005

Ardila, A. (2012). Interaction between lexical and grammatical language systems in the brain. Physics of Life Reviews, 9(2), 198–214. 10.1016/j.plrev.2012.05.001

Berl, M. M., Mayo, J., Parks, E. N., Rosenberger, L. R., VanMeter, J., Ratner, N. B., Vaidya, C. J., & Gaillard, W. D. (2014). Regional differences in the developmental trajectory of lateralization of the language network. Human Brain Mapping, 35(1), 270–284. 10.1002/hbm.22179

Blank, I., Kanwisher, N., & Fedorenko, E. (2014). A functional dissociation between language and multiple-demand systems revealed in patterns of BOLD signal fluctuations. Journal of Neurophysiology, 112(5), 1105–1118. 10.1152/jn.00884.2013

Campbell, K. L., & Tyler, L. K. (2018). Language-related domain-specific and domain-general systems in the human brain. Current Opinion in Behavioral Sciences, 21, 132–137. 10.1016/j.cobeha.2018.04.008

Charbonnier, L., Raemaekers, M. A. H., Cornelisse, P. A., Verwoert, M., Braun, K. P. J., Ramsey, N. F., & Vansteensel, M. J. (2020). A Functional Magnetic Resonance Imaging Approach for Language Laterality Assessment in Young Children. Frontiers in Pediatrics, 8. https://www.frontiersin.org/articles/10.3389/fped.2020.587593

Chater, N., & Christiansen, M. H. (2018). Language acquisition as skill learning. Current Opinion in Behavioral Sciences, 21, 205–208. 10.1016/j.cobeha.2018.04.001

Duncan, J. (2010). The multiple-demand (MD) system of the primate brain: Mental programs for intelligent behaviour. Trends in Cognitive Sciences, 14(4), 172–179. 10.1016/j.tics.2010.01.004

Edgin, J., Clark, C., Massand, E., & Karmiloff-Smith, A. (2015). Building an adaptive brain across development: Targets for neurorehabilitation must begin in infancy. Frontiers in Behavioral Neuroscience, 9. https://www.frontiersin.org/articles/10.3389/fnbeh.2015.00232

Enge, A., Friederici, A. D., & Skeide, M. A. (2020). A meta-analysis of fMRI studies of language comprehension in children. NeuroImage, 215, 116858. 10.1016/j.neuroimage.2020.116858

Fedorenko, E., Behr, M. K., & Kanwisher, N. (2011). Functional specificity for high-level linguistic processing in the human brain. Proceedings of the National Academy of Sciences, 108(39), 16428–16433. 10.1073/pnas.1112937108

Fedorenko, E., Duncan, J., & Kanwisher, N. (2012). Language-Selective and Domain-General Regions Lie Side by Side within Broca’s Area. Current Biology, 22(21), 2059–2062. 10.1016/j.cub.2012.09.011

Fedorenko, E., Duncan, J., & Kanwisher, N. (2013). Broad domain generality in focal regions of frontal and parietal cortex. Proceedings of the National Academy of Sciences of the United States of America, 110(41), 16616–16621. 10.1073/pnas.1315235110

Fedorenko, E., Hsieh, P.-J., Nieto-Castañón, A., Whitfield-Gabrieli, S., & Kanwisher, N. (2010). New Method for fMRI Investigations of Language: Defining ROIs Functionally in Individual Subjects. Journal of Neurophysiology, 104(2), 1177–1194. 10.1152/jn.00032.2010

Friederici, A. D., Brauer, J., & Lohmann, G. (2011). Maturation of the Language Network: From Inter- to Intrahemispheric Connectivities. PLOS ONE, 6(6), e20726. 10.1371/journal.pone.0020726

Gratton, C., Laumann, T. O., Nielsen, A. N., Greene, D. J., Gordon, E. M., Gilmore, A. W., Nelson, S. M., Coalson, R. S., Snyder, A. Z., Schlaggar, B. L., Dosenbach, N. U. F., & Petersen, S. E. (2018). Functional Brain Networks Are Dominated by Stable Group and Individual Factors, Not Cognitive or Daily Variation. Neuron, 98(2), 439–452.e5. 10.1016/j.neuron.2018.03.035

Greve, D. N., & Fischl, B. (2009). Accurate and robust brain image alignment using boundary-based registration. NeuroImage, 48(1), 63–72. 10.1016/j.neuroimage.2009.06.060

Hamrick, P., Lum, J. A. G., & Ullman, M. T. (2018). Child first language and adult second language are both tied to general-purpose learning systems. Proceedings of the National Academy of Sciences, 115(7), 1487–1492. 10.1073/pnas.1713975115

Hannagan, T., Amedi, A., Cohen, L., Dehaene-Lambertz, G., & Dehaene, S. (2015). Origins of the specialization for letters and numbers in ventral occipitotemporal cortex. Trends in Cognitive Sciences, 19(7), 374–382. 10.1016/j.tics.2015.05.006

Holland, S. K., Vannest, J., Mecoli, M., Jacola, L. M., Tillema, J.-M., Karunanayaka, P. R., Schmithorst, V. J., Yuan, W., Plante, E., & Byars, A. W. (2007). Functional MRI of language lateralization during development in children. International Journal of Audiology, 46(9), 533–551. 10.1080/14992020701448994

Holm, S. (1979). A Simple Sequentially Rejective Multiple Test Procedure. Scandinavian Journal of Statistics, 6(2), 65–70.

Juch, H., Zimine, I., Seghier, M. L., Lazeyras, F., & Fasel, J. H. D. (2005). Anatomical variability of the lateral frontal lobe surface: Implication for intersubject variability in language neuroimaging. NeuroImage, 24(2), 504–514. 10.1016/j.neuroimage.2004.08.037

Karmiloff-Smith, A. (2015). An alternative to domain-general or domain-specific frameworks for theorizing about human evolution and ontogenesis. AIMS Neuroscience, 2(2), 91–104. 10.3934/Neuroscience.2015.2.91

Lenneberg, E. H. (1967). The Biological Foundations of Language. Hospital Practice, 2(12), 59–67. 10.1080/21548331.1967.11707799

Li, J., Osher, D. E., Hansen, H. A., & Saygin, Z. M. (2020). Innate connectivity patterns drive the development of the visual word form area. Scientific Reports, 10(1), 18039. 10.1038/s41598-020-75015-7

Lipkin, B., Tuckute, G., Affourtit, J., Small, H., Mineroff, Z., Kean, H., Jouravlev, O., Rakocevic, L., Pritchett, B., Siegelman, M., Hoeflin, C., Pongos, A., Blank, I. A., Struhl, M. K., Ivanova, A., Shannon, S., Sathe, A., Hoffmann, M., Nieto-Castañón, A., & Fedorenko, E. (2022). Probabilistic atlas for the language network based on precision fMRI data from >800 individuals. Scientific Data, 9(1), Article 1. 10.1038/s41597-022-01645-3

Malik-Moraleda, S., Ayyash, D., Gallée, J., Affourtit, J., Hoffmann, M., Mineroff, Z., Jouravlev, O., & Fedorenko, E. (2021). The universal language network: A cross-linguistic investigation spanning 45 languages and 12 language families [Preprint]. Neuroscience. 10.1101/2021.07.28.454040

McCauley, S. M. (2020). Towards an integrated, single-system account of language development as skill learning. Journal of Communication Disorders, 83. 10.1016/j.jcomdis.2019.105942

McMullen, E., & Saffran, J. R. (2004). Music and Language: A Developmental Comparison. Music Perception, 21(3), 289–311. 10.1525/mp.2004.21.3.289

Miller, C. A. (2006). Developmental Relationships Between Language and Theory of Mind. American Journal of Speech-Language Pathology, 15(2), 142–154. 10.1044/1058-0360(2006/014)

Ojemann, G. (1991). Cortical organization of language. The Journal of Neuroscience, 11(8), 2281–2287. 10.1523/JNEUROSCI.11-08-02281.1991

Olulade, O. A., Seydell-Greenwald, A., Chambers, C. E., Turkeltaub, P. E., Dromerick, A. W., Berl, M. M., Gaillard, W. D., & Newport, E. L. (2020). The neural basis of language development: Changes in lateralization over age. Proceedings of the National Academy of Sciences, 117(38), 23477–23483. 10.1073/pnas.1905590117

Rakison, D. H., & Yermolayeva, Y. (2011). How to Identify a Domain-General Learning Mechanism When You See One. Journal of Cognition and Development, 12(2), 134–153. 10.1080/15248372.2010.535228

RStudio Team (2021). RStudio: Integrated Development Environment for R. RStudio, PBC, Boston, MA URL http://www.rstudio.com/.

Saffran, J. R., & Thiessen, E. D. (2007). Domain-general learning capacities. In Blackwell handbook of language development (pp. 68–86). Blackwell Publishing. 10.1002/9780470757833.ch4

Saygin, Z. M., Osher, D. E., Norton, E. S., Youssoufian, D. A., Beach, S. D., Feather, J., Gaab, N., Gabrieli, J. D. E., & Kanwisher, N. (2016). Connectivity precedes function in the development of the visual word form area. Nature Neuroscience, 19(9), 1250–1255. 10.1038/nn.4354

Schettini, E., Hiersche, K. J., & Saygin, Z. M. (2023). Individual Variability in Performance Reflects Selectivity of the Multiple Demand Network among Children and Adults. Journal of Neuroscience, 43(11), 1940–1951. 10.1523/JNEUROSCI.1460-22.2023

Sharp, H., & Hillenbrand, K. (2008). Speech and Language Development and Disorders in Children—ClinicalKey. 55(5), 1159–1173.

Simmons, W. K., Bellgowan, P. S. F., & Martin, A. (2007). Measuring selectivity in fMRI data. Nature Neuroscience, 10(1), Article 1. 10.1038/nn0107-4

Szwed, M., Dehaene, S., Kleinschmidt, A., Eger, E., Valabrègue, R., Amadon, A., & Cohen, L. (2011). Specialization for written words over objects in the visual cortex. NeuroImage, 56(1), 330–344. 10.1016/j.neuroimage.2011.01.073

Tomaiuolo, F., MacDonald, J. D., Caramanos, Z., Posner, G., Chiavaras, M., Evans, A. C., & Petrides, M. (1999). Morphology, morphometry and probability mapping of the pars opercularis of the inferior frontal gyrus: An in vivo MRI analysis. The European Journal of Neuroscience, 11(9), 3033–3046. 10.1046/j.1460-9568.1999.00718.x

Ullman, M. T. (2004). Contributions of memory circuits to language: The declarative/procedural model. Cognition, 92(1), 231–270. 10.1016/j.cognition.2003.10.008

Vigneau, M., Beaucousin, V., Hervé, P. Y., Duffau, H., Crivello, F., Houdé, O., Mazoyer, B., & Tzourio-Mazoyer, N. (2006). Meta-analyzing left hemisphere language areas: Phonology, semantics, and sentence processing. NeuroImage, 30(4), 1414–1432. 10.1016/j.neuroimage.2005.11.002

Wang, J., Yamasaki, B. L., Weiss, Y., & Booth, J. R. (2021). Both frontal and temporal cortex exhibit phonological and semantic specialization during spoken language processing in 7- to 8-year-old children. Human Brain Mapping, 42(11), 3534–3546. 10.1002/hbm.25450

Weiss, Y., Cweigenberg, H. G., & Booth, J. R. (2018). Neural specialization of phonological and semantic processing in young children. Human Brain Mapping, 39(11), 4334–4348. 10.1002/hbm.24274

Weiss-Croft, L., & Baldeweg, T. (2015). Maturation of language networks in children: A systematic review of 22years of functional MRI | Elsevier Enhanced Reader. 123, 269–281. 10.1016/j.neuroimage.2015.07.046

West, G., Shanks, D. R., & Hulme, C. (2021). Sustained Attention, Not Procedural Learning, is a Predictor of Reading, Language and Arithmetic Skills in Children. Scientific Studies of Reading, 25(1), 47–63. 10.1080/10888438.2020.1750618

Xiao, Y., Brauer, J., Lauckner, M., Zhai, H., Jia, F., Margulies, D. S., & Friederici, A. D. (2016). Development of the Intrinsic Language Network in Preschool Children from Ages 3 to 5 Years. PLOS ONE, 11(11), e0165802. 10.1371/journal.pone.0165802

Xiao, Y., Friederici, A. D., Margulies, D. S., & Brauer, J. (2016). Longitudinal changes in resting-state fMRI from age 5 to age 6years covary with language development. NeuroImage, 128, 116–124. 10.1016/j.neuroimage.2015.12.008

Yu, X., Ferradal, S., Sliva, D. D., Dunstan, J., Carruthers, C., Sanfilippo, J., Zuk, J., Zöllei, L., Boyd, E., Gagoski, B., Grant, P. E., & Gaab, N. (2020). Infant functional connectivity fingerprints predict long-term language and pre-literacy outcomes (p. 2020.10.29.360081). bioRxiv. 10.1101/2020.10.29.360081

