## Supplemental Materials for "Functional dissociation of the language network and other cognition in early childhood"

**Supplemental Methods**

**Dilated frontal parcels:**

Prior to CVS registration to subject specific anatomy, the frontal language parcels (from <https://evlab.mit.edu/funcloc/>) were dilated by a factor of three using the Freesurfer’s

mri_binarize command. This decision was made following manual inspection of first-level GLM results, in which the frontal parcels were not covering language activation hotspots. Additionally, the frontal parcels were much smaller than the temporal parcels, and therefore not covering as much activation per subject. Examples from one subject is included in **Supplemental Figure 1** of parcels and activation pre- and post- dilation. Temporal parcels were not dilated, as they covered activation hotspots and were much larger than frontal parcels to begin with.

**Supplemental Results:**

Main results are reported with full cross-sectional sample (N=36) that contain 2 runs of language task and at least one run of the working memory task. However, these subjects moved significantly more on the working memory task (t(35)=2.55, p=0.015, d=0.48); therefore, main results differentiating language and selectivity during the working memory task (selectivity to cognitive control) are presented in a motion matched sample (N=28, t(27)=1.81, p=0.082, d=0.34).

Results in the motion matched sample replicated the dissociation between language selectivity and selectivity to cognitive control in the language network, with significantly higher language selectivity than selectivity to cognitive control. In an ANOVA of task (language and working memory) by fROI (each hemisphere separatel), we found only a significant main effect of task (LH: task: F(1,23)=6.84, p=0.015, η^2^=0.097, nonsignificant: fROI: F(4,92)=1.11, p=0.36, η^2^=0.008; interaction: F(4,92)=1.48, p=0.22, η^2^=0.010; RH: task: F(1,24)=11.63, p=0.002, η^2^=0.121, nonsignificant: fROI: F(4,96)=1.75, p=0.15, η^2^=0.010; interaction: F(4,96)=0.80, p=0.53, η^2^=0.005), such that language selectivity is significantly higher than working memory selectivity across all fROIs (LH: t(23)=2.62, p=0.016, d=0.53; RH: t(24)=3.41, p=0.002, d=0.68).

Additionally, we did not find a relationship between language and selectivity to cognitive cross across ages in our motion matched sample. Language and cognitive control responses within the language fROIs were not correlated (LH: r=-0.037, p=0.85; RH: r=0.24, p=0.22). These findings did not change when controlling for age (partial correlations; LH: r=0.0067, p=0.97; RH: r=0.25, p=0.21).

**Supplemental Figure 1:** Language activation in an example subject with dilated frontal language parcels to show better coverage of hotspots.

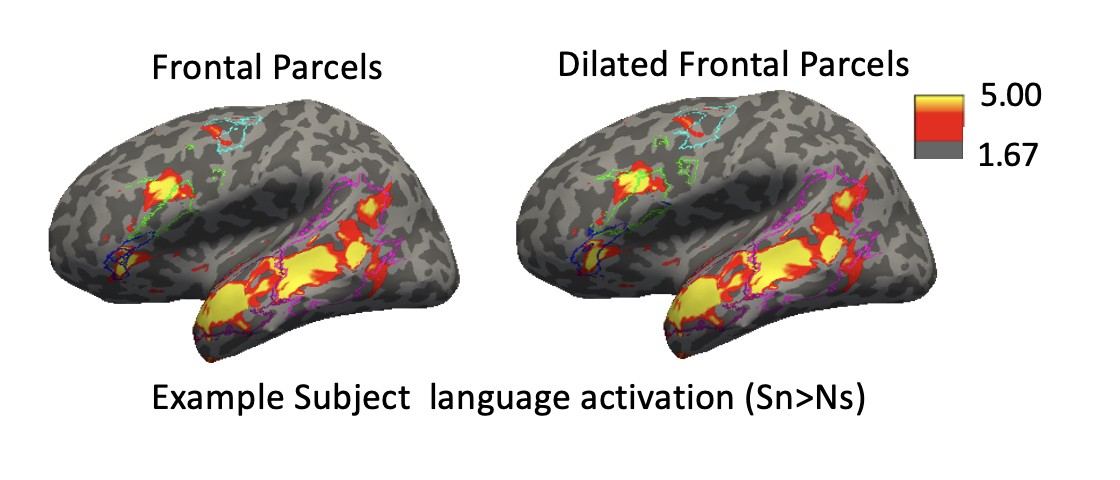

**Supplemental Figure 1:** Language significant activation for an example subject (thresholded at P_uncorrected_ < 0.01) with original (left) and dilated (right) frontal parcels. Middle frontal parcel in teal, Inferior frontal in green, and orbital inferior frontal in dark blue. Temporal parcels were not dilated (posterior temporal in purple, superior temporal in pink).
